# Biomarker-based patient selection improves stroke rehabilitation trial efficiency

**DOI:** 10.1101/459776

**Authors:** Cathy M. Stinear, Winston D. Byblow, P. Alan Barber, Suzanne J. Ackerley, Marie-Claire Smith, Steven C. Cramer

## Abstract

**Background and Purpose:** Inter-subject variability complicates trials of novel stroke rehabilitation therapies, particularly in the sub-acute phase after stroke. We tested whether selecting patients using motor evoked potential (MEP) status, a physiological biomarker of motor system function, could improve trial efficiency.

**Methods:** A retrospective analysis of data from 207 patients (103 women, mean (SD) 70.6 (15.1) years) was used to estimate sample sizes and recruitment rates required to detect a 7-point difference between hypothetical control and treatment groups in upper-limb Fugl-Meyer and Action Research Arm Test scores at 90 days post-stroke. Analyses were carried out for the full sample and for subsets defined by motor evoked potential (MEP) status.

**Results:** Selecting patients according to MEP status reduced the required sample size by 75% compared to an unselected sample. The estimated time needed to recruit the required sample was also reduced by 72% for patients with MEPs, and was increased by 2-3-fold for patients without MEPs.

**Conclusions:** Using biomarkers to select patients can improve stroke rehabilitation trial efficiency by reducing the sample size and recruitment time needed to detect a clinically meaningful effect of the tested intervention.

## Introduction

The acute treatment of ischemic stroke has been transformed by trials that select patients for reperfusion therapies based on advanced imaging biomarkers.^1^ Rehabilitation research could also adopt an approach that uses biomarkers to select patients with the greatest potential to benefit from a putative therapy.^2^ However, stroke is a heterogeneous condition with high inter-individual variability in recovery.^3^ This reduces statistical power in rehabilitation trials, particularly during the sub-acute stage.^2^ A potential biomarker for hand and arm motor recovery is the presence of motor evoked potentials (MEPs) determined using transcranial magnetic stimulation (TMS). Patients with MEPs have a functionally intact motor cortex and lateral corticospinal tract, and better motor recovery.^4^ We determined whether trial efficiency could be improved by using MEP status for patient selection.

## Materials and Methods

The largest available dataset of patients with acute stroke and known baseline MEP status^5^ was used to estimate the sample sizes required to detect rehabilitation benefits on upper-limb motor performance at 90 days after stroke. Inclusion criteria were age ≥18 years and monohemispheric ischemic stroke or intracerebral hemorrhage with new upper-limb weakness within the previous 72 hours. Exclusion criteria were contraindications to TMS, cerebellar stroke, communication or cognitive impairment precluding informed consent, and residing out of area precluding follow-up. Only 2% of screened patients were contraindicated for TMS.^5^ All participants provided written informed consent and completed usual care.

Baseline clinical assessments were made and MEP status of the paretic upper-limb determined within 7 days post-stroke. Participants were considered MEP+ if an MEP of any amplitude and consistent latency (± 3ms) could be elicited in the paretic extensor carpi radialis or first dorsal interosseous muscles on half of at least eight consecutive trials, with the target muscles either at rest or during attempted voluntary activation.^5^ Upper-limb motor impairment was evaluated with the upper-limb Fugl-Meyer scale (UE-FM, maximum=66). Upper-limb motor function was evaluated with the Action Research Arm Test (ARAT, maximum=57). These scales are valid, reliable, and recommended by international consensus;^6^ scores were obtained 90 days after stroke.

Analyses were carried out for the full sample, and repeated for the MEP+ and MEP- subsets. Population estimates of UE-FM and ARAT scores 90 days post-stroke were obtained using bootstrapping with replacement, 10,000 samples, and bias corrected accelerated estimates of 95% confidence intervals. These estimates were then used to calculate the sample sizes required to detect treatment effects of 7 points on the UE-FM and ARAT scores 90 days after stroke, assuming two-tailed tests, alpha=0.05, and 1:1 randomization, with statistical power of 0.80 and 0.90. A 7-point treatment effect was chosen because it is approximately 10% of each scale, and there are currently no agreed upon minimal clinically important differences for these scales at the sub-acute stage of stroke.

## Results

In all, 207 patients (103 women, mean (SD) 70.6 (15.1) years) completed baseline and 90-day follow-up assessments (Table 1). The full sample and MEP+ subset (n=177) had similar demographic and baseline clinical characteristics. The MEP- subset (n=30) had more severe stroke and greater upper limb impairment at baseline. As expected, MEP+ patients had higher estimated population mean 90-day UE-FM and ARAT scores than MEP- patients. The estimated percentage of patients who are MEP+ was 85.5% (95%CI 81.2%-89.9%) giving a relative recruitment rate of 0.855 for MEP+ and 0.145 for MEP-, compared to recruiting patients regardless of MEP status.

**Table 1.** Baseline clinical and demographic characteristics.

|  | Full Sample | MEP+ | MEP- |
| --- | --- | --- | --- |
|  | N = 207 | N = 177 | N = 30 |
| Age (y), mean (range) | 70.6 (18 – 98) | 71.5 (18 – 98) | 65.3 (32 – 90) |
| Sex, female (%) | 103 (49.8) | 84 (47.5) | 19 (61.2) |
| Ethnicity, n (%) |  |  |  |
| European | 131 (63.3) | 117 (66.1) | 14 (46.7) |
| Māori | 10 (4.8) | 9 (5.1) | 1 (3.3) |
| Pacific | 30 (14.5) | 21 (11.9) | 9 (30.0) |
| Asian | 36 (17.4) | 30 (16.9) | 6 (20.0) |
| Stroke risk factors, n (%) |  |  |  |
| Hypertension | 133 (64.5) | 112 (63.3) | 21 (70.0) |
| Dyslipidemia | 66 (31.9) | 53 (29.9) | 13 (43.3) |
| Previous cardiac disease | 56 (27.1) | 52 (29.4) | 4 (13.3) |
| Atrial fibrillation | 47 (22.7) | 41 (23.2) | 6 (20.0) |
| Diabetes mellitus | 43 (20.8) | 35 (19.8) | 8 (26.7) |
| Ex-smoker | 35 (16.9) | 31 (17.5) | 4 (13.3) |
| Smoker | 17 (8.2) | 12 (6.8) | 5 (16.7) |
| First stroke, n (%) | 181 (87.4) | 155 (87.6) | 26 (86.7) |
| Stroke type, n (%) |  |  |  |
| TACI<sup>a</sup> | 12 (5.8) | 6 (3.4) | 6 (20.0) |
| PACI<sup>b</sup> | 74 (35.7) | 59 (33.3) | 15 (50.0) |
| LACI<sup>c</sup> | 84 (40.6) | 78 (44.1) | 6 (20.0) |
| POCI<sup>d</sup> | 16 (7.7) | 14 (7.9) | 2 (6.7) |
| ICH<sup>e</sup> | 21 (10.1) | 20 (11.3) | 1 (3.3) |
| Hemisphere, left, n (%) | 99 (47.8) | 93 (52.5) | 6 (20.0) |
| Hand, dominant, n (%) | 95 (45.9) | 88 (49.7) | 7 (23.3) |
| Intravenous thrombolysis, n (%) | 19 (9.2) | 17 (9.6) | 2 (6.7) |
| Endovascular thrombectomy, n (%) | 3 (1.4) | 1 (0.6) | 2 (6.7) |
| NIHSS<sup>f</sup> score, median (range) | 4 (0 – 19) | 4 (0 – 16) | 11 (2 – 19) |
| NIHSS score 0 – 4, n (%) | 112 (54.1) | 111 (62.7) | 1 (3.3) |
| NIHSS score 5 – 15, n (%) | 85 (41.1) | 62 (35.0) | 23 (76.7) |
| NIHSS score ≥ 16, n (%) | 10 (4.8) | 4 (2.3) | 6 (20.0) |
| UE-FM<sup>g</sup> score |  |  |  |
| Mean (SD) | 41.6 (20.8) | 47.2 (16.9) | 8.5 (3.8) |
| Range | 2 – 65 | 2 – 65 | 4 – 21 |

a. Total anterior circulation infarct.
b. Partial anterior circulation infarct.
c. Lacunar infarct.
d. Posterior circulation infarct, excluding cerebellar stroke.
e. Intracerebral hemorrhage.
f. National Institutes of Health Stroke Scale.
g. Upper-extremity Fugl-Meyer scale, maximum = 66, higher scores indicate less motor impairment.

For both UE-FM and ARAT, the estimated required sample size to detect a treatment effect at 90 days for MEP+ patients was only 27 – 29% compared to the full sample of unselected patients (Table 2). Recruiting the required sample size for MEP+ patients would thus take approximately one third of the time needed to recruit unselected patients. The estimated required samples size to detect a treatment effect at 90 days for MEP- patients was 31% for UE-FM, and 20% for ARAT, relative to the full sample (Table 2). However, recruiting the required sample size for MEP- patients would take approximately three times longer for the UE-FM, and almost twice as long for ARAT, than for unselected patients.

**Table 2.** Estimates of 90 day scores, required sample sizes, and relative time to recruit, with 95% confidence intervals.

|  | Full Sample<br>N = 207 | MEP+<br>N = 177 | MEP-<br>N = 30 |
| --- | --- | --- | --- |
| UE-FM<sup>a</sup> score at 90 days |  |  |  |
| Estimated mean<sup>b</sup> | 52.2 (49.8 – 54.6) | 58.2 (56.8 – 59.5) | 17.1 (13.5 – 21.2) |
| Estimated SD<sup>b</sup> | 17.1 (15.0 – 18.9) | 8.7 (7.3 – 10.1) | 11.3 (8.2 – 13.6) |
| Sample size<sup>c</sup> |  |  |  |
| Statistical power = 0.80 | 190 (148 – 232) | 52 (38 – 68) | 82 (46 – 122) |
| Statistical power = 0.90 | 254 (196 – 319) | 68 (48 – 90) | 112 (60 – 162) |
| Relative time to recruit<sup>d</sup> |  |  |  |
| Statistical power = 0.80 |  | 0.32 (0.30 – 0.34) | 2.98 (2.14 – 3.63) |
| Statistical power = 0.90 |  | 0.31 (0.29 – 0.33) | 3.04 (2.11 – 3.50) |
| ARAT<sup>e</sup> score at 90 days |  |  |  |
| Estimated mean<sup>b</sup> | 43.6 (41.2 – 46.0) | 49.8 (48.3 – 51.2) | 7.2 (4.3 – 10.4) |
| Estimated SD<sup>b</sup> | 17.7 (15.7 – 19.4) | 9.5 (7.9 – 11.0) | 8.8 (6.8 – 10.1) |
| Sample size<sup>c</sup> |  |  |  |
| Statistical power = 0.80 | 204 (160 – 244) | 60 (42 – 80) | 52 (32 – 68) |
| Statistical power = 0.90 | 272 (214 – 316) | 80 (56 – 106) | 70 (42 – 90) |
| Relative time to recruit<sup>d</sup> |  |  |  |
| Statistical power = 0.80 |  | 0.34 (0.31 – 0.38) | 1.76 (1.34 – 1.92) |
| Statistical power = 0.90 |  | 0.34 (0.31 – 0.39) | 1.78 (1.35 – 1.96) |

- Upper-extremity Fugl-Meyer scale, max=66, higher scores indicate less motor impairment. - Estimates of population means and standard deviations (SD) for scores 90 days after stroke, obtained using bootstrapping. - Required sample size to detect a 7 point difference in scores between hypothetical treatment and control groups at 90 days after stroke. - Time to recruit the required sample size of MEP+ or MEP- patients, relative to the time required to recruit the required sample size of unselected patients. - Action Research Arm Test, max=57, higher scores indicate better motor performance.

## Discussion

MEP status is a biologically relevant and non-invasive biomarker of corticospinal tract function that clinicians can obtain at the bedside after minimal training.^5^ Selecting patients on the basis of MEP+ status reduces variance in outcome measures at 90 days, and is estimated to reduce sample size and recruitment time by around two-thirds, without limiting the pool of participants. Selecting patients on the basis of MEP- status also reduces variance and the required sample size by at least two-thirds. Corticospinal tract biomarkers might also be relevant for broader outcomes, such as independence, and warrant further investigation along with neuroimaging and blood biomarkers.

This small retrospective study reflects the characteristics of patients who may be suitable for participation in trials of upper-limb rehabilitation interventions. It serves as an example of how biomarkers could be used to enrich samples for stroke rehabilitation trials. Patients, clinicians, investigators, and funding agencies stand to gain from improved trial efficiency. Using biomarkers for patient selection in stroke rehabilitation trials could markedly increase trial sensitivity and efficiency, with associated decreases in participant burden, research costs, and time required for completion.

## Acknowledgements

We thank Matthew Petoe PhD for assistance with data acquisition, and Sharon Yeatts PhD for advice on statistical analyses.

